# Tetracycline exposure alters key gut microbiota in Africanized honey bees (*Apis mellifera scutellata* x spp.)

**DOI:** 10.1101/2021.05.31.446413

**Authors:** Kilmer Oliveira Soares, Celso José Bruno de Oliveira, Adriana Evangelista Rodrigues, Priscylla Carvalho Vasconcelos, Núbia Michelle Vieira da Silva, Octavio Gomes da Cunha Filho, Christopher Madden, Vanessa L. Hale

## Abstract

Honey bees play a critical role in ecosystem health, biodiversity maintenance, and crop yield. Antimicrobials, such as tetracyclines, are used widely used across agriculture, medicine, and in bee keeping, and bees can be directly or indirectly exposed to tetracycline residues in the environment. In European honey bees, tetracycline exposure has been linked with shifts in the gut microbiota that negatively impact bee health. However, the effects of antimicrobials on Africanized honey bee gut microbiota have not been examined. The aim of this study was to investigate the effects of tetracycline exposure on the gut microbial community of Africanized honey bees (*Apis mellifera scutellata* x spp), which are important pollinators in South, Central, and North America. Bees (n=1,000) were collected from hives in Areia-PB, Northeastern Brazil, placed into plastic chambers and kept under controlled temperature and humidity conditions. The control group (CON) was fed daily with syrup (10g) consisting of a 1:1 solution of demerara sugar and water, plus a solid protein diet (10g) composed of 60% soy extract and 40% sugar syrup. The tetracycline group (TET) was fed identically but with the addition of tetracycline hydrochloride (450 µg/g) to the sugar syrup. Bees were sampled from each group before (day 0), and after tetracycline exposure (days 3, 6 and 9). Abdominal contents dissected out of each bee underwent DNA extraction and 16S rRNA sequencing (V3-V4) on an Illumina MiSeq. Sequences were filtered and processed through QIIME2 and DADA2. Microbial community composition and diversity and differentially abundant taxa were evaluated by treatment and time. Bee gut microbial composition (Jaccard) and diversity (Shannon) differed significantly and increasingly over time and between CON and TET groups. Tetracycline exposure was associated with decreased relative abundances of *Bombella* and *Fructobacillus*, along with decreases in key core microbiota such as *Snodgrassella, Gilliamella*, Rhizobiaceae, and *Apibacter*. These microbes are critical for nutrient metabolism and pathogen defense, and decreased abundances of these microbes could negatively affect bee health. Considering the global ecological and economic importance of honey bees as pollinators, it is critical to understand the effects of agrochemicals including antimicrobials on honey bees.

## Introduction

Bees play a critical role as pollinators in ecosystems across the globe, contributing to the maintenance of biodiversity on Earth (Kevan and Viana, 2003; Michener, 2007). In addition to this important ecological function, bees are also essential as pollinators in agriculture systems (Gisder and Genersch, 2017; Hung et al., 2018). Honey bees (*Apis* spp.), specifically, are the top crop pollinators and directly enhance crop yields (Gisder and Genersch, 2017). The Africanized honey bee (*Apis mellifera scutellata* x spp.), a crossbreed between European honey bees (*Apis mellifera* sspp.) and African honey bees (*Apis mellifera scutellata*), emerged in the late 1950’s in Brazil (Winston, 1992). African honey bees adapted and spread widely across the Americans because of their reproductive traits and superior ability to colonize tropical ecosystems compared with European bees. Some of the traits include improved thermoregulation capacity, greater resistance to diseases, increased egg-laying rates, more frequent queen replacement, and shorter developmental time (Guzmán-Novoa et al., 2011).

In spite of their great economic and biological importance, bee populations across the planet have been under increasing threat due to human population expansion, habitat destruction, and the use of agrochemicals including pesticides and antimicrobials. The use of such compounds has been associated with an increased occurrence of Colony Collapse Disorder (CCD), a phenomenon characterized by the disappearance of worker bees and compromise of the honey bee colony (Caires et al., 2017; Raymann et al., 2017; Motta et al., 2018). Despite potential links between agrochemicals and CCD, agrochemical use, and specifically antimicrobial use in livestock production (Thaker et al., 2010; Park et al., 2017), is projected to increase 67% by 2030, and nearly double in developing countries including Brazil, Russia, India, China, and South Africa (Van Boeckel et al., 2015). According to a recent report on global antimicrobial use in livestock (OIE, 2018), tetracyclines were the most commonly used antimicrobial class among the 116 countries that provided data. Moreover, tetracyclines represented approximately 35% of the antimicrobial use in these countries, including use for growth promotion in feed animals, which is an ongoing practice in many countries. Recently, tetracyclines were also highlighted as an option for the treatment and prophylaxis of COVID-19, and tetracycline use has increased significantly in some hospitals during the pandemic (Sodhi and Etminan, 2020; Peñalva et al., 2021).

Importantly, tetracycline is poorly absorbed by mammalian hosts and 30 to 90% of the drug is excreted in active forms in urine and feces (Khan and Ongerth, 2004; Chee-Sanford et al., 2009; Watkinson et al., 2009). This can result in increased antimicrobial contamination in wastewater and farm runoff (Borrely et al., 2012; Faria et al., 2016; Hendriksen et al., 2019). Tetracycline residues have been detected in irrigation water (0.14 ppm), pig waste lagoons (0.7 ppm), soil (25 ppm), hospital effluents (0.53 ppm), and at wastewater treatment plants (0.92 ppm) (Meyer et al., 2000; Pena et al., 2010; Wang et al., 2014). Although prohibited in Brazil and Europe, oxytetracycline is also used to control bacterial infections in fruit trees including *Candidatus Liberibacter* spp., the causative agent of Citrus Greening Disease (Chanvatik et al., 2019), and *Xylella fastidiosa*, which causes Pierce’s disease in grapevines (Hopkins, 1979). In these cases, oxytetracycline is sprayed over orchards or vineyards, and oxytetracycline concentrations on plant tissues can range from 100 to 4,166 ppm (Chanvatik et al., 2019).

Bees can be indirectly exposed to antimicrobials while foraging in these agricultural or urban environments that contain tetracycline residues (Lau and Nieh, 2016). Bees can also be directly exposed to tetracyclines in the course of treatment for European and American foulbrood, bacterial diseases that cause severe losses in hives and honey production (Doughty et al., 2004; Martel et al., 2006). To treat foulbrood, oxytetracycline is applied directly onto the hives at doses ranging from 500 (Dinkov et al., 2005) to 5900 ppm (Kochansky, 2000). Antimicrobials can disturb gut microbial communities and affect their overall structure and function (Blaser, 2014). Gut microbes are critical to host health (Pessione, 2012; Clark and Mach, 2017; Monda et al., 2017) and play a role in immune system development, biosynthesis of vitamins (LeBlanc et al., 2013) and hormones (Clarke et al., 2014), and cellulose degradation (Warnecke et al., 2007). Antimicrobial-induced alterations in the gut microbiota compromise nutrient metabolism (Lee et al., 2014) and pathogen defense mechanisms in European honey bees (Koch and Schmid-Hempel, 2011; Engel et al., 2012; Martinson et al., 2012; Kwong et al., 2017; Motta et al., 2018; Wang et al., 2021). However, there is no information regarding the effects of antimicrobials on the gut microbiota of Africanized honey bees (Tian et al., 2012; Raymann et al., 2017; Wu et al., 2020).

Considering the widespread prevalence of tetracycline in the environment due to its use in agriculture, medicine, and in relation hive health, and evidence of gut microbiome disturbances in European honey bees due to antimicrobial exposure, the aim of this study was to investigate the effects of tetracycline on the gut microbiota of Africanized honey bees (*Apis mellifera scutellata* x spp.) in tropical conditions.

## Methods

### Experimental design and sampling

The study was carried out in December 2019 at the Bee Laboratory (LABE) of the Federal University of Paraiba, Areia -PB (6° 58’20” S; 35° 43’16.9” W; Altitude 545 m). The average annual temperature of Paraiba is 22.54 ° C; the average relative humidity is 83.65%; and the annual precipitation in 2019 was 1360.2 mm (INMET, 2020).

On Day 0 (D0), approximately one thousand bees were collected from five outdoor hives at LABE, placed into ten plastic chambers that were kept in an incubator at 32°C and 66% relative humidity (TE-371, Tecnal, Piracicaba, Brazil) (**Figure 1**). The plastic chambers measured 176.71 cm^2^ and were covered with a nylon screen. The bees were divided into two groups: The control group (CON), was fed daily with 10g of syrup consisting of a 1:1 solution of demerara sugar and water. Sterile cotton balls were soaked into the syrup and then placed into the bee chambers daily. Bees were also fed a solid protein diet (10g) composed of 60% soy extract and 40% demerara syrup solution. The tetracycline group (TET) was fed identically except that syrup contained 450 µg/g (equivalent to 450 ppm) tetracycline hydrochloride (Tetramed, Medquímica, Brazil). This dose reflects what honey bees may be exposed to within some agricultural environments and is also within the range of hive dosing for the treatment of foulbrood (Raymann et al., 2017).

**Figure 1.**
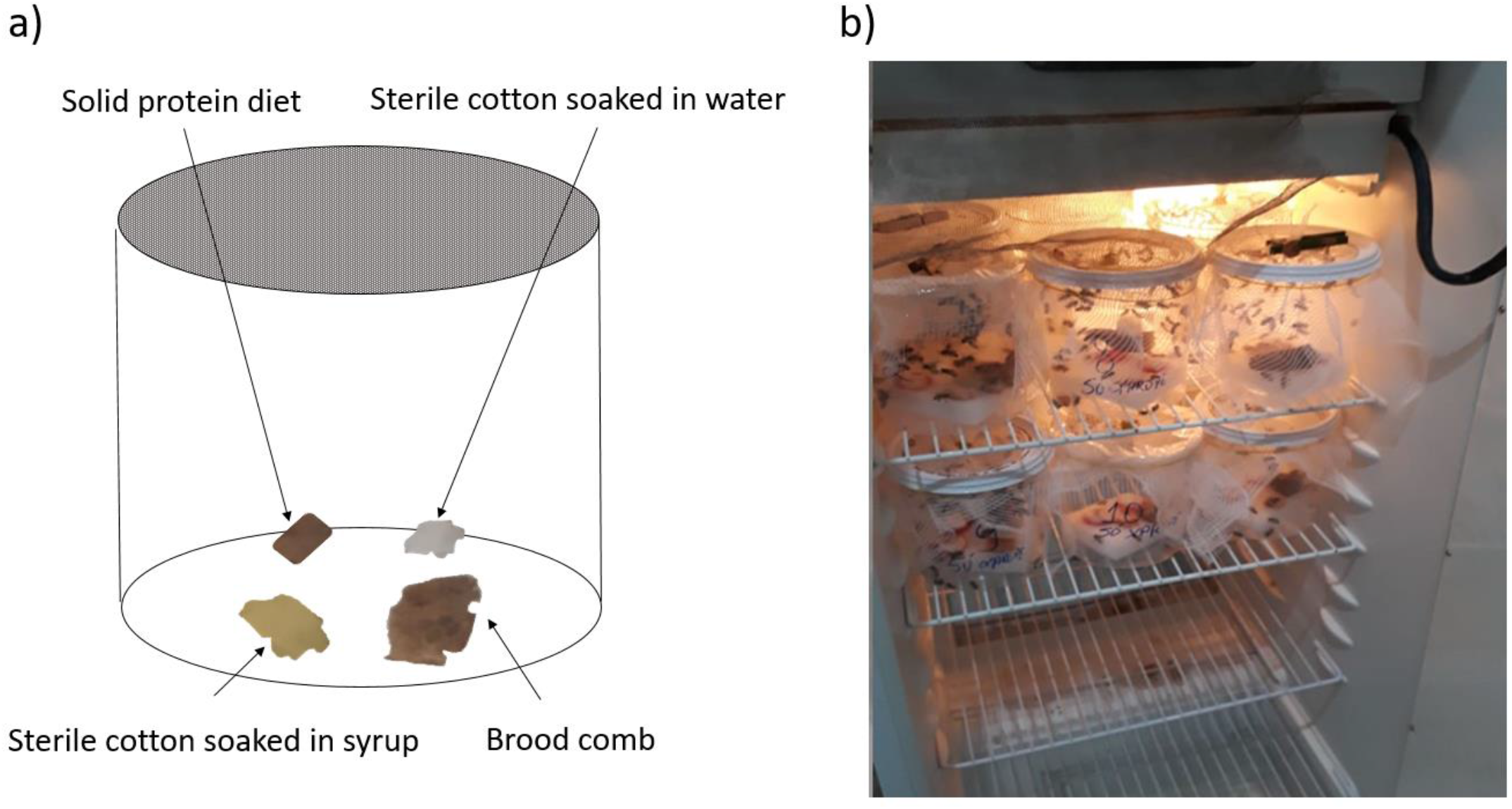
a) Approximately 100 bees were housed in each plastic chamber along with a piece of brood comb. Sterile cotton soaked in water or sugar syrup and a solid protein diet were also included in each chamber, and chambers were covered with Nylon screen. B) All chambers were placed in an incubator that was maintained at 32°C ± 1.45 temp and 66% ± 5.34 relative humidity for the duration of the experiment.

A 9 cm^2^-piece of brood comb was placed in each chamber. Five replicates of 20 bees each were collected from each group at each sampling point including: day 0 (D0, pre-treatment) and days three (D3), six (D6), and nine (D9) (**Figure 2**). Bees were placed in sterile tubes containing 70% alcohol, transported to the lab and stored at -20°C until extraction. All procedures performed were approved by the Biodiversity Authorization and Information System – SISBIO (Protocol #: 71750-1, approved on 09/19/2019).

**Figure 2.**
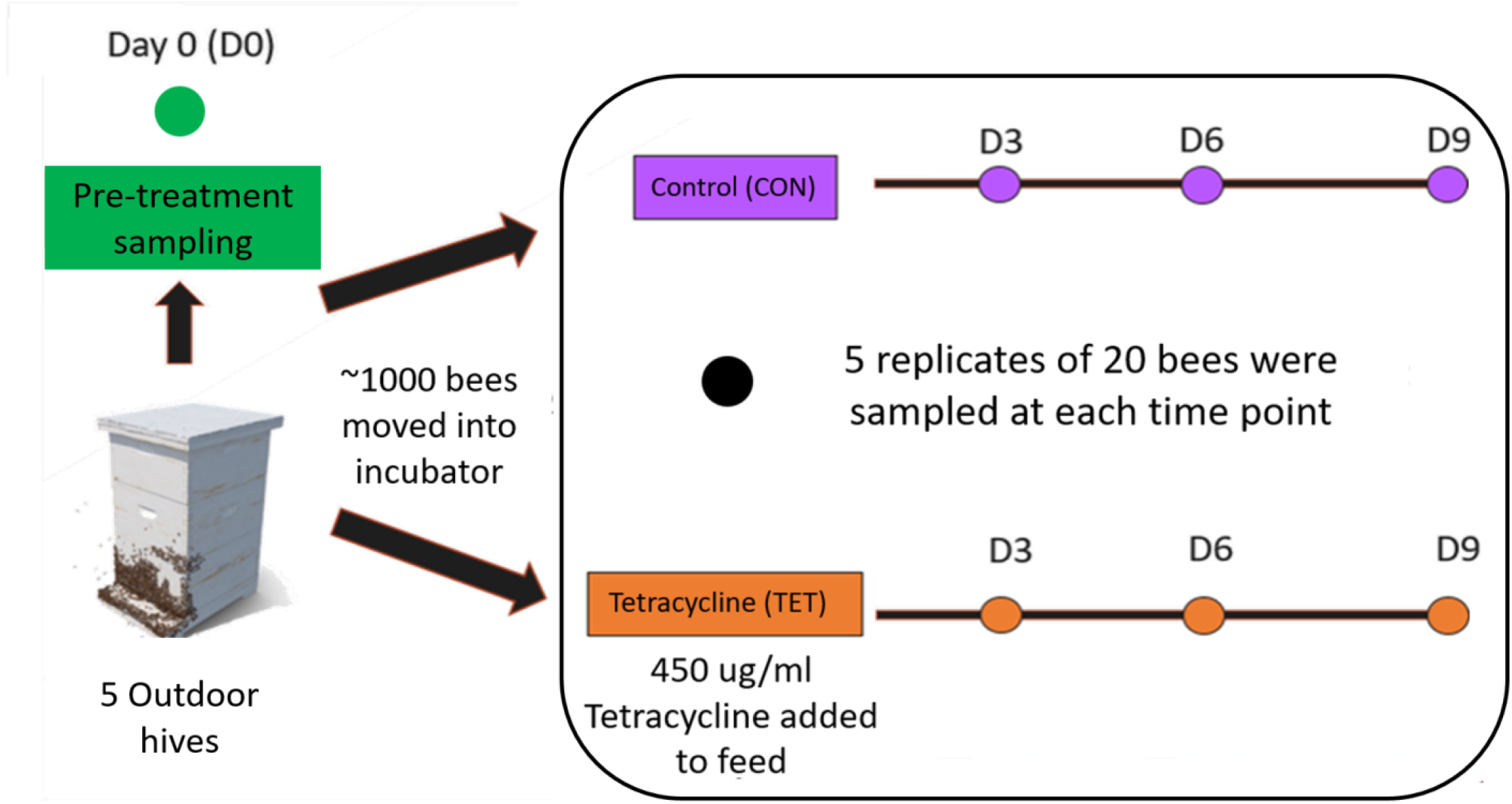
Experimental design. Five replicates of 20 bees were collected from the control (CON) and tetracycline (TET) groups at each time point including Day 0 (D0 – Pre-treatment), and Days 3 (D3), 6 (D6), and 9 (D9).

### DNA extraction, library preparation, and sequencing

Prior to extraction, bees were placed on sterile filter paper for 10 minutes for defrosting and alcohol evaporation. Bee intestines were dissected by using a sterile pair of scissors to make a cross-sectional cut across the last segment of the bee abdomen. With sterile tweezers, abdominal content was collected out of the abdomen and transferred into microtubes. Abdominal contents from 20 bees were pooled into a single tube for DNA extraction, which was performed using a commercial kit (PowerSoil DNA Isolation kit, Qiagen, Germany) following the manufacturer’s protocol. After extraction, DNA was electrophorized in agarose gel for quality analysis and quantified using a microvolume spectrophotometer (Colibri LB 915, Titertek-Berthold, Germany). DNA concentrations were quantified by fluorometry (Qubit 2.0, Life Invitrogen, USA) before further processing steps.

The V3-V4 region of the microbial 16S rRNA gene was amplified by PCR using 2.5 μL template DNA (5 ng/μL), 5 μL forward primer, 5 μL reverse primer, and 12 μL 2X KAPA HiFi HotStart ReadyMix (KAPA Biosystems, Wilmington, MA, USA) in a total volume of 25 μL. The following primers were used: 341F (5′–TCG TCG GCA GCG TCA GAT GTG TAT AAG AGA CAG CCT ACG GGN GGC WGC A–3′) and 805R (5′–GTC TCG TGG GCT CGG AGA TGT GTA TAA GAG ACA GGA CTA CHV GGG TAT CTA ATC C–3′). PCR reaction conditions were as follows: Initial denaturation at 95°C for 3 min, followed by 25 cycles at 95°C for 30 s, 55°C for 30 s, and 72°C for 30s and a final extension to 72°C for 5 min.

Amplification products were visualized in 1.5% agarose gel before purification using magnetic beads (AMPure XP, Beckman Coulter, USA) to remove excess primer. The dual indices and Illumina sequencing adapters were attached using a Nextera XT Index Kit (Illumina). A second clean up step was then performed using magnetic beads. The purified PCR products were quantified by fluorometry (Qubit 2.0, Life Invitrogen). For sequencing, pooled libraries were denatured with NaOH, diluted with hybridization buffer, then heat denatured. Paired-end sequencing was performed on an Illumina MiSeq with a V2 kit (2 ⨯ 250 cycles). At least 5% PhiX DNA was added for sequencing control purposes (Kit PhiX, Illumina). Negative controls including blanks (no template) that underwent the extraction along with all of the other samples and samples from each of the feeds.

### Sequence processing and statistical analyses

The raw demultiplexed paired-end sequences were processed using QIIME 2-2020.2 (Bolyen et al., 2019). Reads were filtered, denoised, and truncated to a length of 248 base pairs, and then parsed for non-chimeric sequences using DADA2, producing Amplicon Sequence Variants (ASV) (Callahan et al., 2016). Sequences were aligned using “qiime fragment-insertion sepp” for phylogenetic analysis (Matsen et al., 2012). Taxonomic composition of the samples were determined rypla pretrained naive Bayes classifier with a 99% sequence similarity threshold for V3-V4 reference sequences (SILVA-132-99-nb-classifier.qza) and the “qiime feature-classifier classify-sklearn”. Negative control samples were examined for potential contaminant taxa. No taxa overlapped between negative control and true samples. Microbial diversity was quantified using Pielou’s (evenness) and Shannon (richness and abundance) diversity indices. ANOVAs were used to compare diversity between groups in R 4.1.0 (Ripley, 2001) after testing for normality using a Shapiro-Wilk test.

Beta diversity was evaluated using Bray-Curtis and Jaccard distances in QIIME 2-2020.2 (Bolyen et al., 2019). Microbial community composition was evaluated by Principle Coordinate Analysis (PCoA) and visualized using the Emperor plugin 2020.2.0 (Vázquez-Baeza et al., 2017). PERMANOVAs were employed as recommended (Anderson, 2001) to test for differences in microbial composition between experimental groups (Pre-treatment vs. CON vs. TET) and over time (Day 0 – pre-treatment, and Days 3, 6, 9).

Differentially abundant taxa between groups were identified using an analysis of composition of microbiomes (ANCOM) (Mandal et al., 2015). We also performed a core microbiota analysis in QIIME2, to identify taxa present in 95% of the samples. The relative abundances of core microbes were then compared by treatment and time using two-way ANOVAa after testing for normality using a Shapiro-Wilk test. A *P*-value < 0.05 was used in the statistical tests for significance.

## Results

### 16S rRNA sequencing reads

We obtained a total of 3,575,254 raw reads across all samples, ranging from 10,268 to 459,284 reads per sample and averaging 102,150 reads per sample. After the denoising process, 3,346,889 (93.61%) were retained for downstream analyses. Reads were classified into 2,140 features which were aligned to 131 different taxa. Reads identified as chloroplasts, mitochondria, unassigned and eukaryota were removed from all samples.

### Microbial composition and diversity in tetracycline-treated bees

Bee gut microbial composition was significantly altered by treatment (pre-treatment, control, tetracycline) (PERMANOVA: Jaccard R2= 0.115, *p* = 0.001) and time (D0, D3, D6, D9) (PERMANOVA: Jaccard R2= 0.046, *p* = 0.001; **Fig. 3, Fig. S1**). Notably, the interaction of treatment and time was also significant (Adonis: R2= 0.035, p-value= 0.024), and the effect of treatment increased over time (**Fig. 3, Fig. S1**). Microbial diversity also differed significantly by time but not by treatment (Two-way ANOVA: Shannon Index treatment *p* = 0.295, time *p* = 0.042; **Fig. 4a**). No pairwise comparisons were significant; although, microbial diversity differences on Day 9 (*p* = 0.081) were greater than at previous timepoints, with the tetracycline group having lower diversity than the control group. This suggests that diversity was being impacted gradually after continual exposure to tetracycline. Microbial community evenness (Pielou’s Index) did not differ significantly by treatment or time (Two-way ANOVA: Pielou’s Index treatment *p* = 0.457, time *p* = 0.061; **Fig. 4b**)

**Figure 3.**
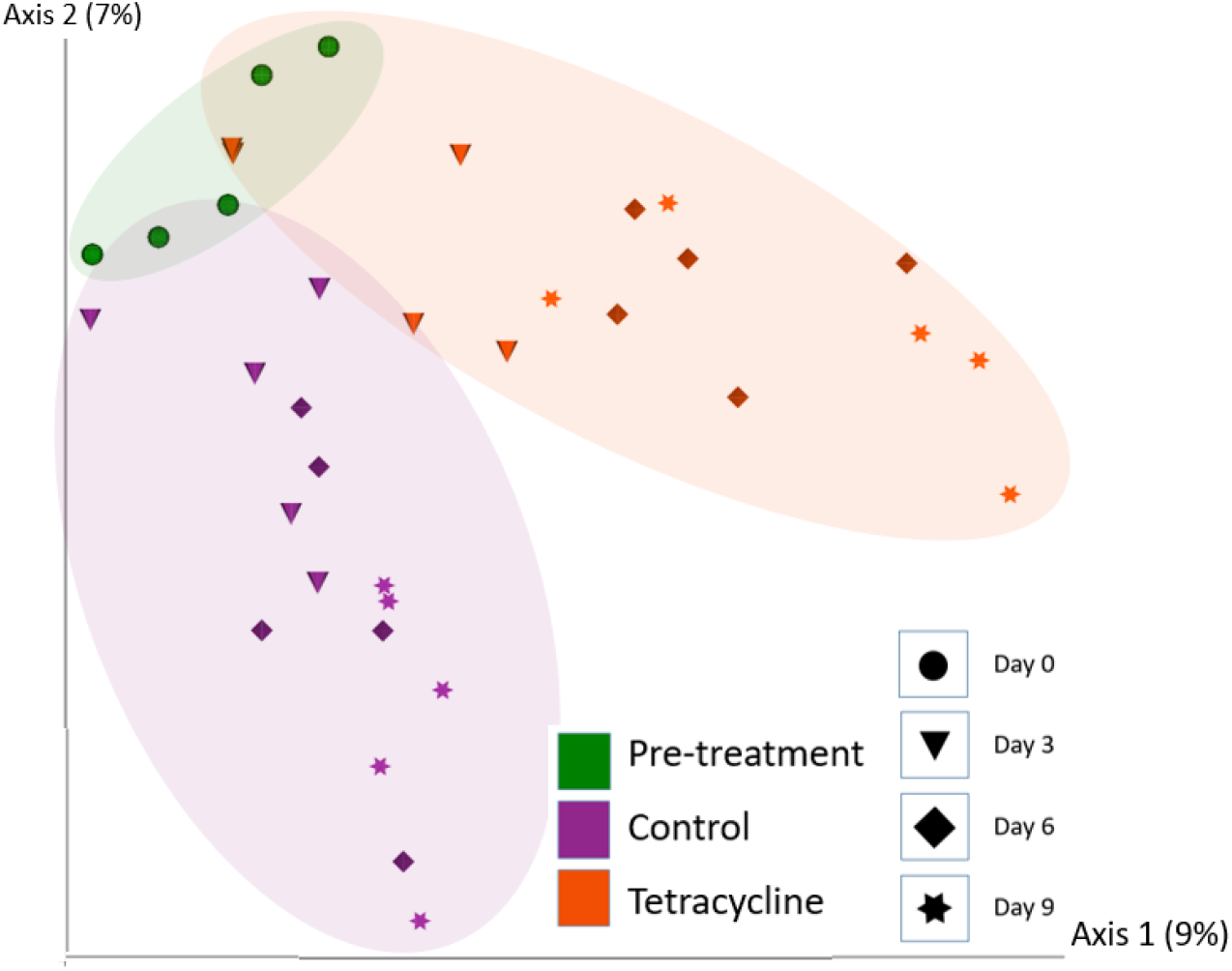
Bee gut microbial composition (Jaccard) based on treatment (Pre-treatment, Control, Tetracycline) and time (Day 0 – pre-treatment, Days 3, 6, 9). Microbial composition was significantly altered by treatment (PERMANOVA: *p* = 0.001) and time (PERMANOVA: *p* = 0.001; also **Supp. Fig. 1**)

**Figure 4.**
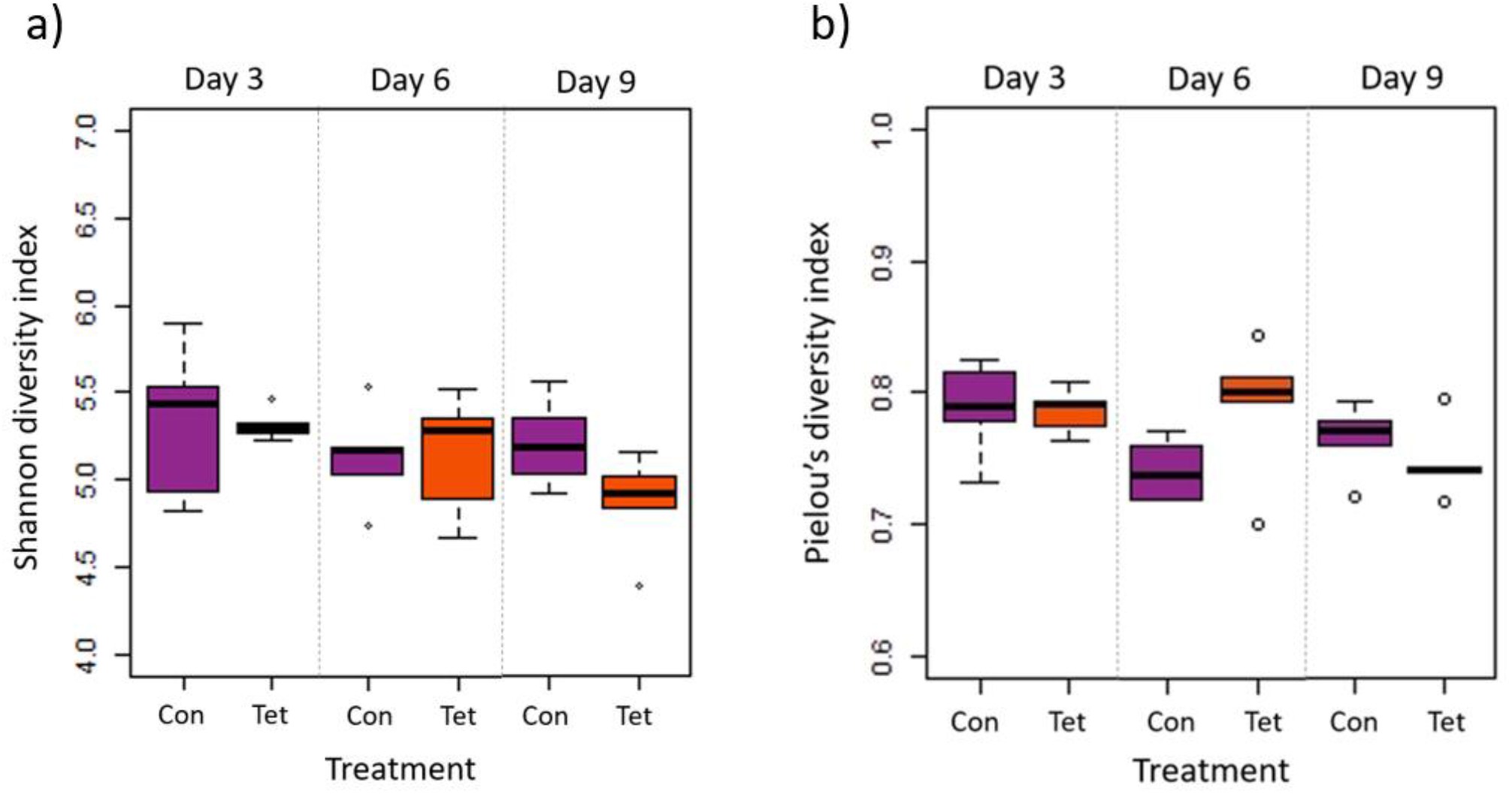
Microbial diversity and evenness by treatment and time. Box plot shows outliers, first and third quartiles (lower and upper edges), and highest, lowest and median values (horizontal black dash) for Control (Con) and Tetracycline (Tet) groups. a) There were significant differences in diversity (Shannon Index) by time (*p* = 0.042) but not treatment (*p* = 0.295); although, no pairwise comparisons were significant. b) There were no significant differences in evenness (Pielou’s Index) by time (*p* = 0.061) or by treatment (*p* = 0.457).

### Core microbiota and differentially abundant taxa

A core microbiota analyses identified eight genera that were present in 95% of the samples across all treatments and times including: *Lactobacillus*, a taxon from the class Gammaproteobacteria, *Bifidobacterium, Snodgrassella, Gilliamella*, a taxon from the family Rhizobiaceae, *Apibacter*, and *Commensalibacter* (**Fig. 5**). These taxa accounted for 22% of all genera in the dataset. We then used a two-way ANOVA to compare relative abundances of these taxa by treatment and time.

**Figure 5.**
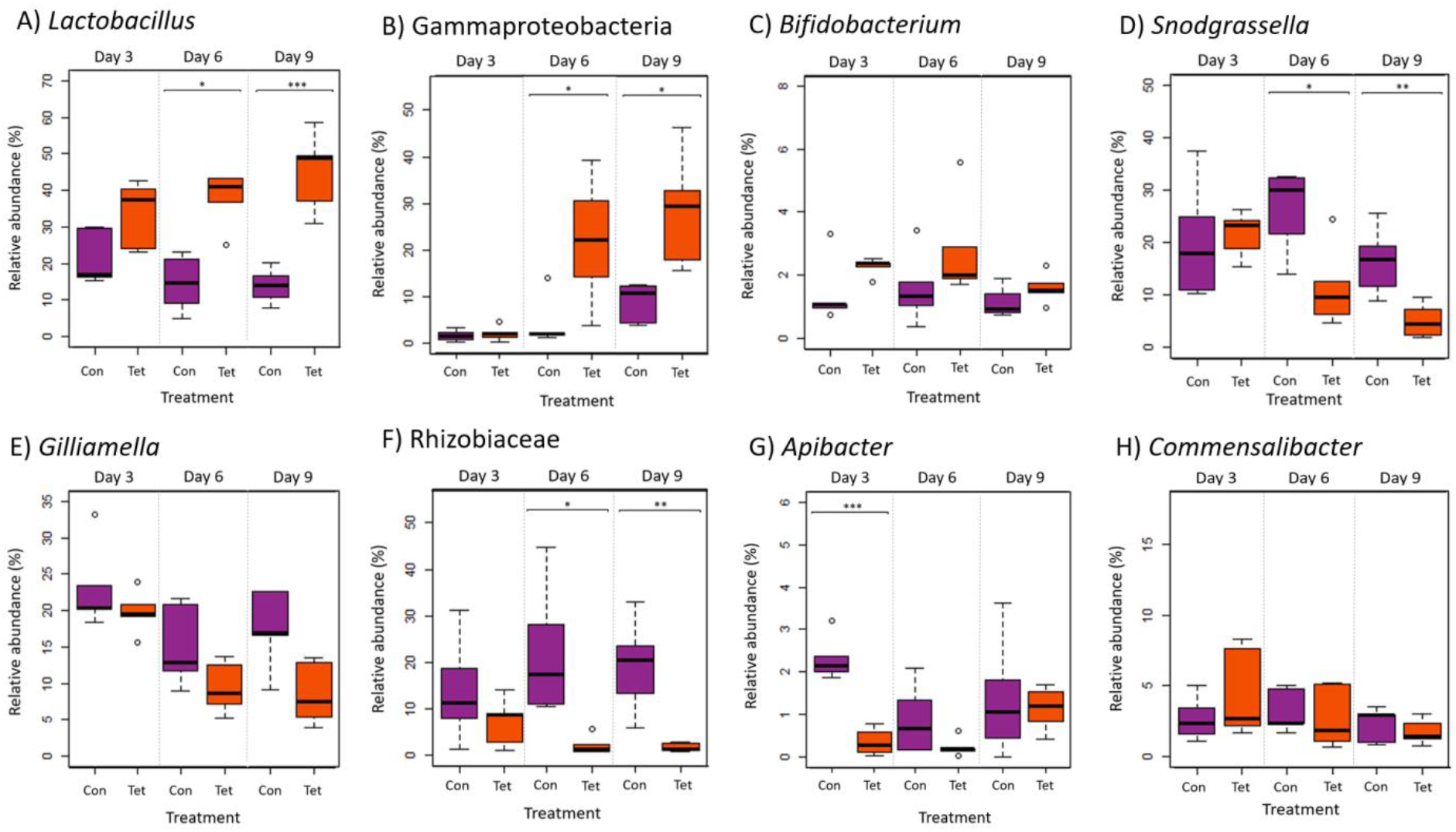
Relative abundances of core microbiota (genera) that were present in 95% of all samples: A) *Lactobacillus*, B) Gammaproteobacteria, C) *Bifidobacterium*, D) *Snodgrassella*, E) *Gilliamella*, F) one taxon from the family Rhizobiaceae, G) *Apibacter* and H) *Commensalibacter*. Box plot shows outliers, first and third quartiles (lower and upper edges), and highest, lowest and median values (horizontal black dash) for Control (Con) and Tetracycline (Tet) groups. (ANOVA: \**p* < 0.05; ** *p* < 0.01 and *** *p* < 0.001)

*Lactobacillus* and the Gammaproteobacteria taxa abundances increased in the tetracycline group over time (Two way ANOVA: *Lactobacillus* treatment *p* < 0.0001, time *p* = 0.684, interaction *p* = 0.049; Gammaproteobacteria treatment *p =* 0.0003, time *p =* 0.0001, interaction Gammaproteobacteria *p =* 0.01; **Fig. 5a,b**). *Bifidobacterium* was also increased in the tetracycline group (*p =* 0.029); although, abundances did not change over time (**Fig. 5c**). Abundances of *Snodgrassella, Gilliamella* and a taxon from the Rhizobiaceae family all decreased over time in the tetracycline group (*Snodgrassella* treatment *p =* 0.007, time *p =* 0.006; *Gilliamella* treatment *p =* 0.01, time *p =* 0.065; Rhizobiaceae treatment *p <* 0.0001, time *p =* 0.98; **Fig. 5d,e,f**). *Apibacter* was also significantly decreased in the tetracycline group; although only at the early time points (treatment *p* < 0.004; time *p* = 0.834; interaction *p* = 0.004; **Fig. 5g**). There were no significant differences in the relative abundances of *Commensalibacter* between groups or over time (p > 0.05; **Fig. 5h**).

An ANCOM identified five differentially abundant taxa by treatment at the L6 genera level including *Bombella* and *Fructobacillus*, a taxon in the family Enterobacteriaceae, *Idiomarina*, and taxon in the class Gammaproteobacteria (**Fig. 6, 5b**). The relative abundances of *Bombella, Fructobacillus*, and the Enterobacteriaceae family taxa differed significantly by treatment (Two-way ANOVA: *Bombella p =* 0.000986; *Fructobacillus p =* 0.0002; Enterobacteriaceae taxa *p* < 0.0001) but not by time (*Bombella p =* 0.115; *Fructobacillus p =* 0.107; Enterobacteriaceae taxa *p =* 0.186), and were decreased in the tetracycline group at all time points (**Fig. 6a,b,c**). *Idiomarina* differed significantly by treatment (Idiomarina *p =* 0.0002) and by time (*Idiomarina p* < 0.0001;), and there was a significant interaction between treatment and time (*Idiomarina p =* 0.005;) as both *Idiomarina* and the Gammaproteobacteria taxa increased over time particularly in the tetracycline group (**Fig. 6d, 5b**). We also performed an ANCOM analysis at the L7 (roughly species) and amplicon sequencing variant levels and produced similar results in terms of differentially abundant microbes: *Fructobacillus* (W=30) and *Bombella* (W=30) species were decreased in the tetracycline group, while 2 Gammaproteobacteria ASVs (W=194, W=179) increased over time in the tetracycline group.

**Figure 6.**
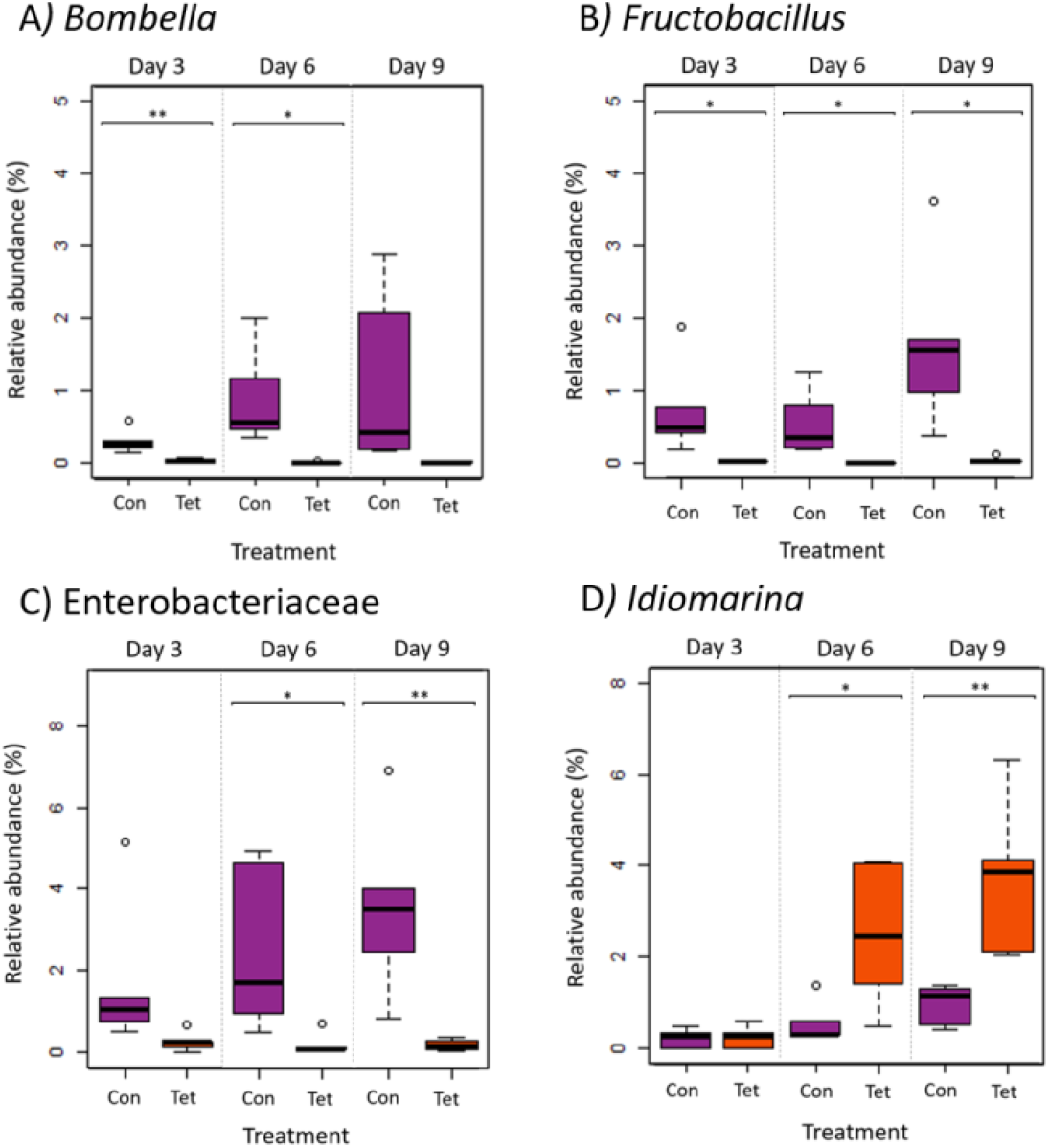
Relative abundances of differentially abundant genera (ANCOM) by treatment and by time. A) *Bombella*, B) *Fructobacillus*, C) a taxon in the family Enterobacteriaceae and D) *Idiomarina*. A Gammaproteobacteria taxa was also identified as a core microbe and a differentially abundant microbe between Con and Tet groups and is shown in **Fig. 5b**. Box plot shows outliers, first and third quartiles (lower and upper edges), and highest, lowest and median values (horizontal black dash) for Control (Con) and Tetracycline (Tet) groups. (ANOVA: * *p* < 0.05; ** p < 0.01 and *** significant at *p* < 0.001)

## Discussion

Our results demonstrated that tetracycline exposure was associated with alterations in Africanized honey bee gut microbial composition but not diversity over time. We further identified shifts in core and non-core microbiota by treatment and time. These tetracycline-linked gut microbial changes suggest negative implications for honey bee nutrient metabolism and pathogen resistance.

### Core microbial taxa and tetracycline treatment

All 8 core microbial taxa identified in this this study (*Lactobacillus*, a taxon of the class Gammaproteobacteria, *Bifidobacterium, Snodgrassella, Gilliamella*, a taxon of the family Rhizobiaceae, *Apibacter*, and *Commensalibacter*) have been previously reported as core microbiota in European honey bees (Engel et al., 2012; Powell et al., 2014; Kwong and Moran, 2016; Raymann et al., 2017; Motta et al., 2018). The increased relative abundance of three core microbes - *Lactobacillus, Bifidobacterium*, and a taxon of the Gammaproteobacteria class – in bees exposed to tetracycline has also been observed in previous studies on bees exposed to chemical compounds or in compromised hives. For instance, increased relative abundances of *Lactobacillus* were reported in bees exposed to the herbicide glyphosate, as well as in hives showing colony collapse disorder (CCD) (Cornman et al., 2012; Motta et al., 2018). Bees in CCD-affected hives also showed increased abundances of Gammaproteobacteria (Cornman et al., 2012), while increased abundances of Bifidobacteriaceae have been reported in bees exposed to coumaphos, an organophosphate insecticide used to control ectoparasites on cattle (Bleau et al., 2020). Taken together, these results suggest that *Lactobacillus, Bifidobacterium*, and Gammaproteobacteria may be positively associated with exposure to agrochemicals. Notably, our results differ from a study on European honey bees that reported decreases in several *Lactobacillius* ASVs following exposure to oxytetracycline (Daisley et al., 2020).

While *Lactobacillus, Bifidobacterium*, and Gammaproteobacteria increased in response to tetracycline exposure, four core taxa decreased in relative abundance under the same treatment: *Snodgrassella, Gilliamella, Apibacter*, and a taxon of the Rhizobiaceae family. These results corroborate previous studies on European bees showing reduced abundances of *Gilliamella, Snodgrassella*, and Rhizobiaceae (specifically *Bartonella apis*) after exposure to tetracycline or glyphosate (Raymann et al., 2017; Motta et al., 2018). Decreased abundances of *B. apis* have also been observed in bees from collapsing colonies (Raymann and Moran, 2018). *Snodgrassella* and *Gillamella* synergistically produce a biofilm on the gut wall (Raymann and Moran, 2018) that serves as barrier against pathogen colonization and translocation (Engel et al., 2012; Martinson et al., 2012; Motta et al., 2018). Moreover, *Snodgrassella* plays an important role in digestion and energy production through the oxidation of fermentation products. *Gilliamella* is involved in nutrient metabolism and is the major degrader of monosaccharides, pectin, and hemicellulose in the bee gut (Engel et al., 2012; Fouad et al., 2016; Zheng et al., 2019). Pectin-rich pollen is large part of the honey bee diet, but bees do not produce pectinases and must rely on gut microbes like *Gilliamella* for pectin metabolism. Like *Snodgrassella* and *Gilliamella, Apibacter* also colonizes the gut wall (Kwong et al., 2018), and some strains of *Apibacter* encode a type VI secretion system (T6SS) (Kwong et al., 2018), which promotes colonization resistance through the delivery of toxic antibacterial proteins into neighboring cells (Steele et al., 2017). Decreased abundances of *Snodgrassella, Gilliamella*, and *Apibacter* could impact nutrient metabolism and pathogen defense in Africanized honey bees.

Differing sensitivity to tetracycline could explain the taxonomic shifts we observed with tetracycline exposure. Gram positive and gram negative bee gut bacteria reportedly have different sensitivities to host-produced antimicrobial peptides including apidaecin and hymenoptaecin. In a previous study by Kwong et al. (2017), gram-positive species (*Lactobacillus* Firm-5, *Bifidobacterium* sp.) were highly resistant to apidaecin and hymenoptaecin, while gram-negative species, particularly *Snodgrassella alvi*, were more sensitive to hymenoptaecin. It is possible that gram positive bacteria, such as *Lactobacillus* and *Bifidobacterium*, which dominate the hindgut, are less sensitive to tetracycline, while gram negative bacteria – such as *Snodgrassella, Gilliamella, Apibacter*, and *Rhizobiaceae*, which are more common in the ileum – are more sensitive to tetracycline and therefore decreased in abundance following tetracycline exposure while *Lactobacillus* and *Bifidobacterium* increased (Powell et al., 2014; Kwong and Moran, 2016; Kešnerová et al., 2020). *Commensalibacter* was the only core microbe that did not vary in relative abundance after tetracycline exposure; however, these bacteria do vary by season and age in honey bees (Ellegaard and Engel, 2019). In sum, alterations in the core microbiota following tetracycline exposure, and particularly decreased abundances of *Snodgrassella, Gilliamella*, Rhizobiaceae, and *Apibacter*, suggest a reduced capacity for pathogen defense and nutrient metabolism which could potentially increase the susceptibility of Africanized honey bees to parasites or infections.

### Differentially abundant microbes by treatment

Among the five differentially abundant taxa identified between treatment groups, three (*Bombella, Fructobacillus*, an Enterobacteriaceae taxon) were decreased in abundance in bees exposed to tetracycline, while two were increased (*Idiomarina*, and a Gammaproteobacteria taxon, which was also identified as a core bacteria). *Bombella*, formerly *Parasaccharibacter apium* (Smith et al., 2020), is positively associated with bee larval development and protection against *Nosema apis* infection (Corby-Harris et al., 2016; Miller et al., 2020). A previous study also showed that exposure to thiacloprid (insecticide) led to *Bombella* reductions in a dose-dependent manner (Liu et al., 2020). The decreased abundance of *Fructobacillus* observed in our study was expected, as these bacteria are known to be highly sensitive to tetracycline (Rokop et al., 2015). *Fructobacillus* is found throughout bee hives (Endo et al., 2011) and it colonizes brood cells, bee bread, and nectar, creating a niche that promotes the growth and inoculation of core microbes into larvae and developing worker bees (Rokop et al., 2015). As such, decreased abundances of *Bombella* and *Fructobacillus* due to tetracycline exposure could negatively affect Africanized honey bee larval development.

To our knowledge, this is the first study characterizing the gut microbiota of Africanized honey bees in relation to tetracycline exposure. However, this study had several limitations. While the microbial shifts we observed suggest negative implications for bee health, we do not have associated immunological, behavioral, fitness, or production data to explicitly support these implications. Secondly, in this study, we selected a tetracycline concentration consistent with that reported in some agricultural or hive applications. However, quantifying the concentration of tetracycline to which bees are actually exposed under natural conditions is challenging and likely highly variable. Third, we observed a shift in the gut microbiota between pre-treatment and *both* the CON and TET groups, suggesting either an age or “incubator effect” due to an altered diet and environment; although, we attempted to replicate natural temperature and humidity conditions as closely as possible within the incubator. Despite this, there were still clear differences between the CON and TET groups over time. Finally, the function of some of the differentially abundant microbes we identified, such as *Apibacter*, have yet to be elucidated. As such, deeper sequencing and associated studies with metabolomics or transcriptomics are necessary to clarify the role of these microbes in the bee gut.

## Conclusion

Tetracycline exposure altered gut microbial composition in Africanized honey bees (*Apis mellifera scutellata* x spp), and was specifically associated with decreased abundances of *Bombella, Fructobacillus, Snodgrassella, Gilliamella*, Rhizobiaceae, and *Apibacter*. These microbes play a key role in nutrient metabolism and pathogen defense, and reduced abundances of these microbes could potentially have negative impacts on bee health. Considering the global ecological and economic importance of honey bees as pollinators, it is critical to understand the effects of antimicrobials widely used across agriculture, medicine, and in bee keeping, on honey bee health, as bees can be directly or indirectly exposed to these drugs in many of their foraging environments. Future studies assessing bee fitness, behavior, immune response, and disease susceptibility in relation to agrochemical exposure will further elucidate the impacts of these gut microbial changes. Understanding how chemicals, like antimicrobials, affect bees is essential to guide agricultural practices that effectively support ecosystem health.

## Additional Requirements

All experimental procedures were previously approved by the Biodiversity Authorization and Information System – SISBIO (Protocol #: 71750-1, approved on 09/19/2019).

## Author Contributions

KOS (conceptualization, methodology, formal analysis, data curation, writing); CJBO (conceptualization, methodology, writing, supervision, resources, funding acquisition); Adriana Evangelista Rodrigues (conceptualization, methodology, resources); PCV (methodology, investigation); NMVS (methodology, investigation), OGCF (methodology, investigation, writing); CM: (software, methodology, formal analysis, resources, supervision, data curation, writing), VLH (conceptualization, software, methodology, formal analysis, resources, supervision, funding, data curation, writing).

## Acknowledgments

The authors are thankful to Leticia Nascimento, Bruno Domingos, and Roberto from the Bee Laboratory (LABE/CCA/UFPB), Thamara Rocha, Wydemberg Araújo, and Elma Leite from the Laboratory for Assessment of Animal-derived Foods (LAPOA/CCA/UFPB), and Morgan V. Evans and Ryan Mrofchak from Hale Laboratory (VPM/OSU), for their valuable help during experiment, sample processing and analyses.

## Reference styles

We used Vancouver system for in-text citations.

## Supplementary material

Figure S1

## Data Availability Statement

Sequencing data is available at NCBI (PRJNA732391). (These data will be made publicly available at publication.)

## Contribution to the Field Statement

Pollination by honey bees is critical for ecosystem health and crop production around the globe. However, bee populations have been threatened by anthropogenic activities worldwide. Antimicrobial drugs, such as tetracycline, are widely used in agriculture and medicine, and bees can be directly or indirect exposed to residues of these drugs in the environment. Previous studies have shown that antimicrobial exposure can cause gut microbial disturbances and affect the health of European honey bees. This study reports the effects of tetracycline exposure on the gut microbiota of Africanized honey bees, which are important pollinators across South, Central, and North America.

## Supplementary material

**Figure S1.**
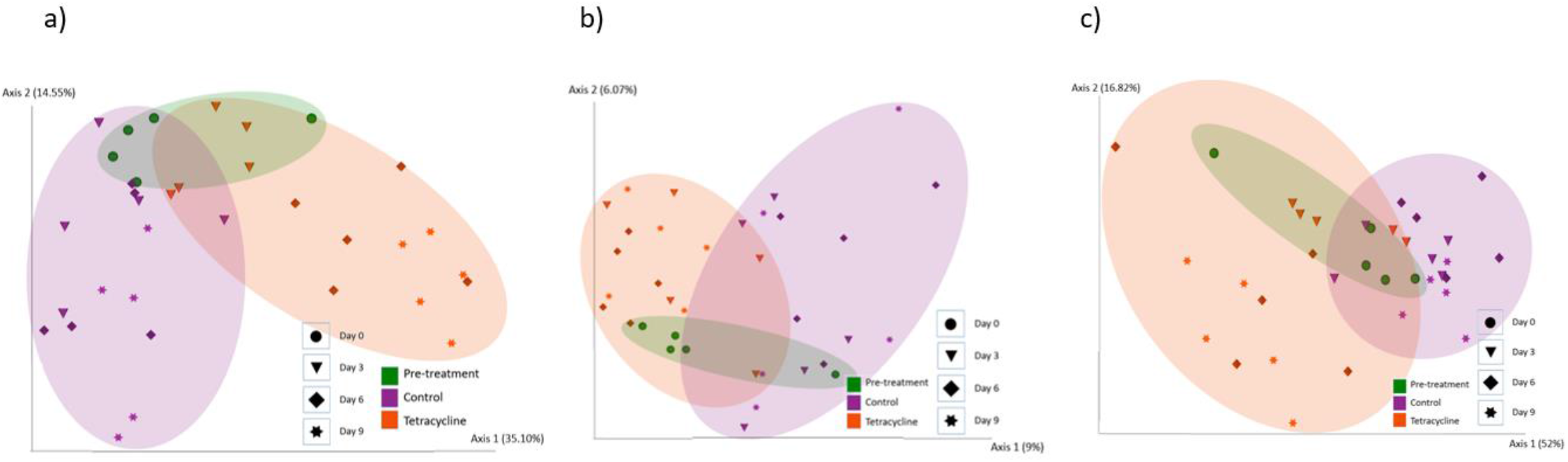
Bee gut microbial composition by treatment (Pre-treatment, Control, Tetracycline) and time (Day 0 – pre-treatment, Days 3, 6, 9) by a) Bray-Curtis, b) Unweighted UniFrac and c) Weighted UniFrac. Microbial composition was significantly altered by treatment (PERMANOVA: Bray Curtis *p* = 0.001; Unweighted UniFrac *p =* 0.001; Weighted UniFrac. *p =* 0.001) and by time (PERMANOVA: Bray-Curtis *p* = 0.001; Unweighted UniFrac *p =* 0.376; Weighted UniFrac *p =* 0.013). Also see **Fig. 3**.

## References

Anderson, M. J. (2001). A new method for non-parametric multivariate analysis of variance. Austral Ecol. 26, 32–46. doi:https://doi.org/10.1111/j.1442-9993.2001.01070.pp.x.

Blaser, M. J. (2014). The microbiome revolution. J. Clin. Invest. 124, 4162–4165. doi:10.1172/JCI78366.

Bleau, N., Bouslama, S., Giovenazzo, P., and Derome, N. (2020). Dynamics of the Honeybee (Apis mellifera) Gut Microbiota Throughout the Overwintering Period in Canada. Microorganisms 8. doi:10.3390/microorganisms8081146.

Bolyen, E., Rideout, J. R., Dillon, M. R., Bokulich, N. A., Abnet, C. C., Al-Ghalith, G. A., et al. (2019). Reproducible, interactive, scalable and extensible microbiome data science using QIIME 2. Nat. Biotechnol. 37, 852–857. doi:10.1038/s41587-019-0209-9.

Borrely, S. I., Caminada, S. M. L., Ponezi, A. N., Santos, D. R. dos, and Silva, V. H. O. (2012). Contaminação das águas por resíduos de medicamentos: ênfase ao cloridrato de fluoxetina. Mundo da Saúde, 556–563.

Caires, S. C., Barcelos, D., and others (2017). Collapse of bees: possible causes and consequences of their disappearance in nature. ACTA Apic. Bras. 5.

Callahan, B. J., McMurdie, P. J., Rosen, M. J., Han, A. W., Johnson, A. J. A., and Holmes, S. P. (2016). DADA2: high-resolution sample inference from Illumina amplicon data. Nat. Methods 13, 581. doi:10.1038/nmeth.3869.

Chanvatik, S., Donnua, S., Lekagul, A., Kaewkhankhaeng, W., Vongmongkol, V., Athipunyakom, P., et al. (2019). Antibiotic use in mandarin production (Citrus reticulata Blanco) in major mandarin-producing areas in Thailand: A survey assessment. PLoS One 14, e0225172. Available at: https://doi.org/10.1371/journal.pone.0225172.

Chee-Sanford, J. C., Mackie, R. I., Koike, S., Krapac, I. G., Lin, Y.-F., Yannarell, A. C., et al. (2009). Fate and transport of antibiotic residues and antibiotic resistance genes following land application of manure waste. J. Environ. Qual. 38, 1086–1108. doi:10.2134/jeq2008.0128.

Clark, A., and Mach, N. (2017). The Crosstalk between the Gut Microbiota and Mitochondria during Exercise. Front. Physiol. 8, 319. doi:10.3389/fphys.2017.00319.

Clarke, G., Stilling, R. M., Kennedy, P. J., Stanton, C., Cryan, J. F., and Dinan, T. G. (2014). Minireview: Gut microbiota: the neglected endocrine organ. Mol. Endocrinol. 28, 1221–1238. doi:10.1210/me.2014-1108.

Corby-Harris, V., Snyder, L., Meador, C. A. D., Naldo, R., Mott, B., and Anderson, K. E. (2016). Parasaccharibacter apium, gen. nov., sp. nov., Improves Honey Bee (Hymenoptera: Apidae) Resistance to Nosema. J. Econ. Entomol. 109, 537–543. doi:10.1093/jee/tow012.

Cornman, R. S., Tarpy, D. R., Chen, Y., Jeffreys, L., Lopez, D., Pettis, J. S., et al. (2012). Pathogen Webs in Collapsing Honey Bee Colonies. PLoS One 7, e43562.

Daisley, B. A., Pitek, A. P., Chmiel, J. A., Gibbons, S., Chernyshova, A. M., Al, K. F., et al. (2020). Lactobacillus spp. attenuate antibiotic-induced immune and microbiota dysregulation in honey bees. Commun. Biol. 3, 534. doi:10.1038/s42003-020-01259-8.

Dinkov, D., Kanelov, I., and Zhelyazkova, I. (2005). Persistence of tetracycline and oxytetracycline in bee honey after improper application on bee families. Bulg. J. Vet. Med 8, 205–509.

Doughty, S., Luck, J., and Goodman, R. (2004). Evaluating alternative antibiotics for control of European Foulbrood disease. Australian Government - Rural Industries Research and Development Corporation.

Ellegaard, K. M., and Engel, P. (2019). Genomic diversity landscape of the honey bee gut microbiota. Nat. Commun. 10, 446. doi:10.1038/s41467-019-08303-0.

Endo, A., Irisawa, T., Futagawa-Endo, Y., Sonomoto, K., Itoh, K., Takano, K., et al. (2011). Fructobacillus tropaeoli sp. nov., a fructophilic lactic acid bacterium isolated from a flower. Int. J. Syst. Evol. Microbiol. 61, 898–902. doi:10.1099/ijs.0.023838-0.

Engel, P., Martinson, V. G., and Moran, N. A. (2012). Functional diversity within the simple gut microbiota of the honey bee. Proc. Natl. Acad. Sci. 109, 11002–11007. doi:10.1073/pnas.1202970109.

Faria, A. C. S. de Godoy, I. de, Sanches, A. A. A., Iglesias, G. A., Candido, S. L., Paz, R. C. R. da, et al. (2016). Detection of resistance genes and evaluation of water quality at zoo lakes in Brazil. Ciência Rural 46, 860–866.

Fouad, A. M., Chen, W., Ruan, D., Wang, S., Xia, W. G., and Zheng, C. T. (2016). Impact of heat stress on meat, egg quality, immunity and fertility in poultry and nutritional factors that overcome these effects: A review. Int. J. Poult. Sci. 15, 81–95. doi:10.3923/ijps.2016.81.95.

Gisder, S., and Genersch, E. (2017). Viruses of commercialized insect pollinators. J. Invertebr. Pathol. 147, 51–59. doi:https://doi.org/10.1016/j.jip.2016.07.010.

Guzmán-Novoa, E., Benítez, A. C., Montaño, L. G. E., and Novoa, G. G. (2011). Colonización, impacto y control de las abejas melíferas africanizadas en México. Vet. México 42, 149–178.

Hendriksen, R. S., Munk, P., Njage, P., van Bunnik, B., McNally, L., Lukjancenko, O., et al. (2019). Global monitoring of antimicrobial resistance based on metagenomics analyses of urban sewage. Nat. Commun. 10, 1124. doi:10.1038/s41467-019-08853-3.

Hopkins, D. L. (1979). Effect of tetracycline antibjotics on pierce ‘ s disease of grapevine in florida. Florida State Hortic. Soc. 92, 284–285.

Hung, K.-L. J., Kingston, J. M., Albrecht, M., Holway, D. A., and Kohn, J. R. (2018). The worldwide importance of honey bees as pollinators in natural habitats. Proc. R. Soc. B Biol. Sci. 285, 20172140. doi:10.1098/rspb.2017.2140.

INMET (2020). Meteorological Database for Teaching and Research 2019.

Kešnerová, L., Emery, O., Troilo, M., Liberti, J., Erkosar, B., and Engel, P. (2020). Gut microbiota structure differs between honeybees in winter and summer. ISME J. 14, 801–814. doi:10.1038/s41396-019-0568-8.

Kevan, P. G., and Viana, B. F. (2003). The global decline of pollination services. Biodiversity 4, 3–8. doi:10.1080/14888386.2003.9712703.

Khan, S. J., and Ongerth, J. E. (2004). Modelling of pharmaceutical residues in Australian sewage by quantities of use and fugacity calculations. Chemosphere 54, 355–367. doi:https://doi.org/10.1016/j.chemosphere.2003.07.001.

Koch, H., and Schmid-Hempel, P. (2011). Socially transmitted gut microbiota protect bumble bees against an intestinal parasite. Proc. Natl. Acad. Sci. 108, 19288 LP – 19292. doi:10.1073/pnas.1110474108.

Kochansky, J. (2000). Analysis of oxytetracycline in extender patties. Apidologie 31, 517–524. doi:10.1051/apido:2000103.

Kwong, W. K., Mancenido, A. L., and Moran, N. A. (2017). Immune system stimulation by the native gut microbiota of honey bees. R. Soc. Open Sci. 4, 170003. doi:10.1098/rsos.170003.

Kwong, W. K., and Moran, N. A. (2016). Gut microbial communities of social bees. Nat. Rev. Microbiol. 14, 374.

Kwong, W. K., Steele, M. I., and Moran, N. A. (2018). Genome Sequences of Apibacter spp., Gut Symbionts of Asian Honey Bees. Genome Biol. Evol. 10, 1174–1179. doi:10.1093/gbe/evy076.

Lau, P., and Nieh, J. (2016). Salt preferences of honey bee water foragers. J. Exp. Biol. 219, 790–796. doi:10.1242/jeb.132019.

LeBlanc, J. G., Milani, C., de Giori, G. S., Sesma, F., van Sinderen, D., and Ventura, M. (2013). Bacteria as vitamin suppliers to their host: a gut microbiota perspective. Curr. Opin. Biotechnol. 24, 160– 168. doi:https://doi.org/10.1016/j.copbio.2012.08.005.

Lee, F. J., Rusch, D. B., Stewart, F. J., Mattila, H. R., and Newton, I. L. G. (2014). Saccharide breakdown and fermentation by the honey bee gut microbiome. Environ. Microbiol. doi:10.1111/1462-2920.12526.

Liu, Y.-J., Qiao, N.-H., Diao, Q.-Y., Jing, Z., Vukanti, R., Dai, P.-L., et al. (2020). Thiacloprid exposure perturbs the gut microbiota and reduces the survival status in honeybees. J. Hazard. Mater. 389, 121818. doi:10.1016/j.jhazmat.2019.121818.

Mandal, S., Van Treuren, W., White, R. A., Eggesbø, M., Knight, R., and Peddada, S. D. (2015). Analysis of composition of microbiomes: a novel method for studying microbial composition. Microb. Ecol. Health Dis. 26, 27663. doi:10.3402/mehd.v26.27663.

Martel, A.-C., Zeggane, S., Drajnudel, P., Faucon, J.-P., and Aubert, M. (2006). Tetracycline residues in honey after hive treatment. Food Addit. Contam. 23, 265–273. doi:10.1080/02652030500469048.

Martinson, V. G., Moy, J., and Moran, N. A. (2012). Establishment of Characteristic Gut Bacteria during Development of the Honeybee Worker. Appl. Environ. Microbiol. 78, 2830 LP – 2840. doi:10.1128/AEM.07810-11.

Matsen, F. A., Hoffman, N. G., Gallagher, A., and Stamatakis, A. (2012). A Format for Phylogenetic Placements. PLoS One 7, 1–4. doi:10.1371/journal.pone.0031009.

Meyer, M. T., Bumgarner, J. E., Varns, J. L., Daughtridge, J. V, Thurman, E. M., and Hostetler, K. A. (2000). Use of radioimmunoassay as a screen for antibiotics in confined animal feeding operations and confirmation by liquid chromatography/mass spectrometry. Sci. Total Environ. 248, 181–187. doi:https://doi.org/10.1016/S0048-9697(99)00541-0.

Michener, C. D. (2007). The Bees of the World. 2nd ed. Baltimore, Maryland: The Johns Hopkins University Press.

Miller, D. L., Smith, E. A., and Newton, I. L. G. (2020). A bacterial symbiont protects honey bees from fungal disease. bioRxiv, 2020.01.21.914325. doi:10.1101/2020.01.21.914325.

Monda, V., Villano, I., Messina, A., Valenzano, A., Esposito, T., Moscatelli, F., et al. (2017). Exercise Modifies the Gut Microbiota with Positive Health Effects. Oxid. Med. Cell. Longev. 2017, 3831972. doi:10.1155/2017/3831972.

Morris, D. E., Cleary, D. W., and Clarke, S. C. (2017). Secondary bacterial infections associated with influenza pandemics. Front. Microbiol. 8, 1041.

Motta, E. V. S., Raymann, K., and Moran, N. A. (2018). Glyphosate perturbs the gut microbiota of honey bees. PNAS 115, 10305–10310. doi:10.1073/pnas.1803880115.

OIE (2018). OIE annual report on antimicrobial agents intended for use in animals. better understanding of the global situation. third annual report.

Park, J., Gasparrini, A. J., Reck, M. R., Symister, C. T., Elliott, J. L., Vogel, J. P., et al. (2017). Plasticity, dynamics, and inhibition of emerging tetracycline resistance enzymes. Nat. Chem. Biol. 13, 730– 736. doi:10.1038/nchembio.2376.

Pena, A., Paulo, M., Silva, L. J. G., Seifrtová, M., Lino, C. M., and Solich, P. (2010). Tetracycline antibiotics in hospital and municipal wastewaters: A pilot study in Portugal. Anal. Bioanal. Chem. 396, 2929–2936. doi:10.1007/s00216-010-3581-3.

Peñalva, G., Benavente, R. S., Pérez-Moreno, M. A., Pérez-Pacheco, M. D., Pérez-Milena, A., Murcia, J., et al. (2021). Effect of the coronavirus disease 2019 pandemic on antibiotic use in primary care. Clin. Microbiol. Infect. doi:10.1016/j.cmi.2021.01.021.

Pessione, E. (2012). Lactic acid bacteria contribution to gut microbiota complexity: lights and shadows. Front. Cell. Infect. Microbiol. 2, 86. doi:10.3389/fcimb.2012.00086.

Powell, J. E., Martinson, V. G., Urban-Mead, K., and Moran, N. A. (2014). Routes of Acquisition of the Gut Microbiota of the Honey Bee (Apis mellifera). Appl. Environ. Microbiol. 80, 7378–7387. doi:10.1128/AEM.01861-14.

Raymann, K., and Moran, N. A. (2018). The role of the gut microbiome in health and disease of adult honey bee workers. Curr. Opin. Insect Sci. 26, 97–104. doi:10.1016/J.COIS.2018.02.012.

Raymann, K., Shaffer, Z., and Moran, N. A. (2017). Antibiotic exposure perturbs the gut microbiota and elevates mortality in honeybees. PLoS Biol. 15, e2001861–e2001861. doi:10.1371/journal.pbio.2001861.

Ripley, B. D. (2001). The R project in statistical computing. MSOR Connect., 23–25. doi:10.11120/msor.2001.01010023.

Rokop, Z. P., Horton, M. A., and Newton, I. L. G. (2015). Interactions between Cooccurring Lactic Acid Bacteria in Honey Bee Hives. Appl. Environ. Microbiol. 81, 7261–7270. doi:10.1128/AEM.01259-15.

Smith, E. A., Anderson, K. E., Corby-Harris, V., McFrederick, Q. S., and Newton, I. L. G. (2020). Reclassification of seven honey bee symbiont strains as Bombella apis. bioRxiv, 2020.05.06.081802. doi:10.1101/2020.05.06.081802.

Sodhi, M., and Etminan, M. (2020). Therapeutic Potential for Tetracyclines in the Treatment of COVID-19. Pharmacother. J. Hum. Pharmacol. Drug Ther. 40, 487–488. doi:doi:10.1002/phar.2395.

Steele, M. I., Kwong, W. K., Whiteley, M., and Moran, N. A. (2017). Diversification of Type VI Secretion System Toxins Reveals Ancient Antagonism among Bee Gut Microbes. MBio 8. doi:10.1128/mBio.01630-17.

Thaker, M., Spanogiannopoulos, P., and Wright, G. D. (2010). The tetracycline resistome. Cell. Mol. Life Sci. 67, 419–431. doi:10.1007/s00018-009-0172-6.

Tian, B., Fadhil, N. H., Powell, J. E., Kwong, W. K., and Moran, N. A. (2012). Long-Term Exposure to Antibiotics Has Caused Accumulation of Resistance Determinants in the Gut Microbiota of Honeybees. MBio 3, e00377–12. doi:10.1128/mBio.00377-12.

Van Boeckel, T. P., Brower, C., Gilbert, M., Grenfell, B. T., Levin, S. A., Robinson, T. P., et al. (2015). Global trends in antimicrobial use in food animals. Proc. Natl. Acad. Sci. 112, 5649 LP – 5654. doi:10.1073/pnas.1503141112.

Vázquez-Baeza, Y., Gonzalez, A., Smarr, L., McDonald, D., Morton, J. T., Navas-Molina, J. A., et al. (2017). Bringing the Dynamic Microbiome to Life with Animations. Cell Host Microbe 21, 7–10. doi:10.1016/j.chom.2016.12.009.

Wang, F.-H., Qiao, M., Lv, Z.-E., Guo, G.-X., Jia, Y., Su, Y.-H., et al. (2014). Impact of reclaimed water irrigation on antibiotic resistance in public parks, Beijing, China. Environ. Pollut. 184, 247–253. doi:https://doi.org/10.1016/j.envpol.2013.08.038.

Wang, K., Li, J., Zhao, L., Mu, X., Wang, C., Wang, M., et al. (2021). Gut microbiota protects honey bees (Apis mellifera L.) against polystyrene microplastics exposure risks. J. Hazard. Mater. 402, 123828. doi:https://doi.org/10.1016/j.jhazmat.2020.123828.

Warnecke, F., Luginbühl, P., Ivanova, N., Ghassemian, M., Richardson, T. H., Stege, J. T., et al. (2007). Metagenomic and functional analysis of hindgut microbiota of a wood-feeding higher termite. Nature 450, 560.

Watkinson, A. J., Murby, E. J., Kolpin, D. W., and Costanzo, S. D. (2009). The occurrence of antibiotics in an urban watershed: From wastewater to drinking water. Sci. Total Environ. 407, 2711–2723. doi:10.1016/j.scitotenv.2008.11.059.

Winston, M. L. (1992). The Biology and Management of Africanized Honey Bees. Annu. Rev. Entomol. 37, 173–193. doi:10.1146/annurev.en.37.010192.001133.

Wu, Y., Zheng, Y., Chen, Y., Wang, S., Chen, Y., Hu, F., et al. (2020). Honey bee (Apis mellifera) gut microbiota promotes host endogenous detoxification capability via regulation of P450 gene expression in the digestive tract. Microb. Biotechnol. 13, 1201–1212. doi:https://doi.org/10.1111/1751-7915.13579.

Zheng, H., Perreau, J., Powell J. E., Han, B., Zhang, Z., Kwong W. K., et al. (2019). Division of labor in honey bee gut microbiota for plant polysaccharide digestion. Proc. Natl. Acad. Sci. 116, 25909 LP – 25916. doi:10.1073/pnas.1916224116.

